# Allosteric Motions of the CRISPR-Cas9 HNH Nuclease Probed by NMR and Molecular Dynamics

**DOI:** 10.1101/660613

**Authors:** Kyle W. East, Jocelyn C. Newton, Uriel N. Morzan, Atanu Acharya, Erin Skeens, Gerwald Jogl, Victor S. Batista, Giulia Palermo, George P. Lisi

## Abstract

CRISPR-Cas9 is a widely employed genome-editing tool with functionality reliant on the ability of the Cas9 endonuclease to introduce site-specific breaks in double-stranded DNA. In this system, an intriguing allosteric communication has been suggested to control its DNA cleavage activity through flexibility of the catalytic HNH domain. Here, solution NMR experiments and a novel Gaussian accelerated Molecular Dynamics (GaMD) simulation method are used to capture the structural and dynamic determinants of allosteric signaling within the HNH domain. We reveal the existence of a millisecond timescale dynamic pathway that spans HNH from the region interfacing the adjacent RuvC nuclease and propagates up to the DNA recognition lobe in full-length CRISPR-Cas9. These findings reveal a potential route of signal transduction within the CRISPR-Cas9 HNH nuclease, advancing our understanding of the allosteric pathway of activation. Further, considering the role of allosteric signaling in the specificity of CRISPR-Cas9, this work poses the mechanistic basis for novel engineering efforts aimed at improving its genome editing capability.

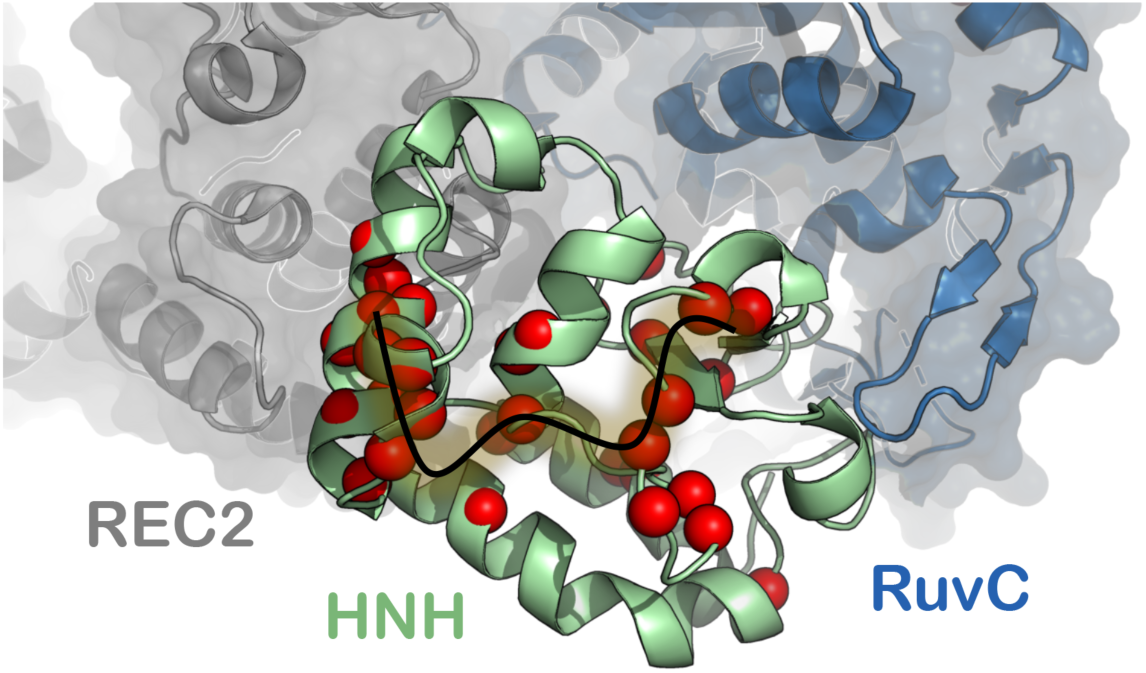

---

The CRISPR-Cas9 enzyme machine has exciting applications in genome editing and numerous investigations have sought to harness its mechanism for therapeutic bioengineering.^1-2^ Cas9 is an RNA-guided DNA endonuclease, which generates double-stranded breaks in DNA by first recognizing its protospacer-adjacent motif (PAM) sequence and then cleaving the two DNA strands via the HNH and RuvC nuclease domains.^3^ Structural studies of Cas9 have employed crystallographic^4-6^ and cryo-EM^7-8^ techniques, revealing several well-defined subdomains, including the catalytic domains, a recognition (REC) lobe and a PAM interacting (PI) region (Figure 1A). In parallel, Förster Resonance Energy Transfer (FRET) techniques provided insight into the large-scale conformational changes that occur during nucleic acid processing.^9-11^ These and other biophysical studies have been invaluable to our current understanding of Cas9 function.^12-13^ Building on this experimental information, computational investigations have been carried out to describe the conformational and dynamic requirements underlying Cas9 mechanistic action. All-atom Molecular Dynamics (MD) simulations have described the conformational activation of the Cas9 protein toward the binding and enzymatic processing of nucleic acids.^14-16^ These investigations also revealed the ability of Cas9 to propagate the DNA binding signal across the HNH and RuvC nuclease domains for concerted cleavage of double-stranded DNA.^17^ Notably, biochemical experiments and MD simulations have jointly indicated a dynamically driven allosteric signal throughout Cas9, where the intrinsic flexibility of the catalytic HNH domain regulates the conformational activation of both nucleases, therefore controlling the DNA cleavage activity ^9, 17^ Detailed knowledge of this allosteric mechanism and of the conformational control exerted by HNH is essential for understanding Cas9 function and for engineering efforts aimed at improving the specificity of this system through modulation of its allosteric signaling.^18^ In this respect, an indepth investigation necessitates the use of experimental techniques such as solution nuclear magnetic resonance (NMR) to quantify the motional timescales critical to this allosteric cross-talk. NMR can readily detect subtle conformational fluctuations at the molecular level, with precise information about the local dynamics on picosecond (ps) to nanosecond (ns) timescales (i.e. the so-called fast dynamics), as well as those occurring over microseconds (µs) to milliseconds (ms) (i.e. slow dynamics), both of which contribute to allosteric signaling.^19-22^ The power of solution NMR is magnified when coupled to MD simulations^23-24^ that capture protein fluctuations and conformations on the same timescales of NMR experiments, offering an interpretation at the atomic scale while also describing the subtle changes that characterize protein allostery.^25-27^

**Figure 1.**
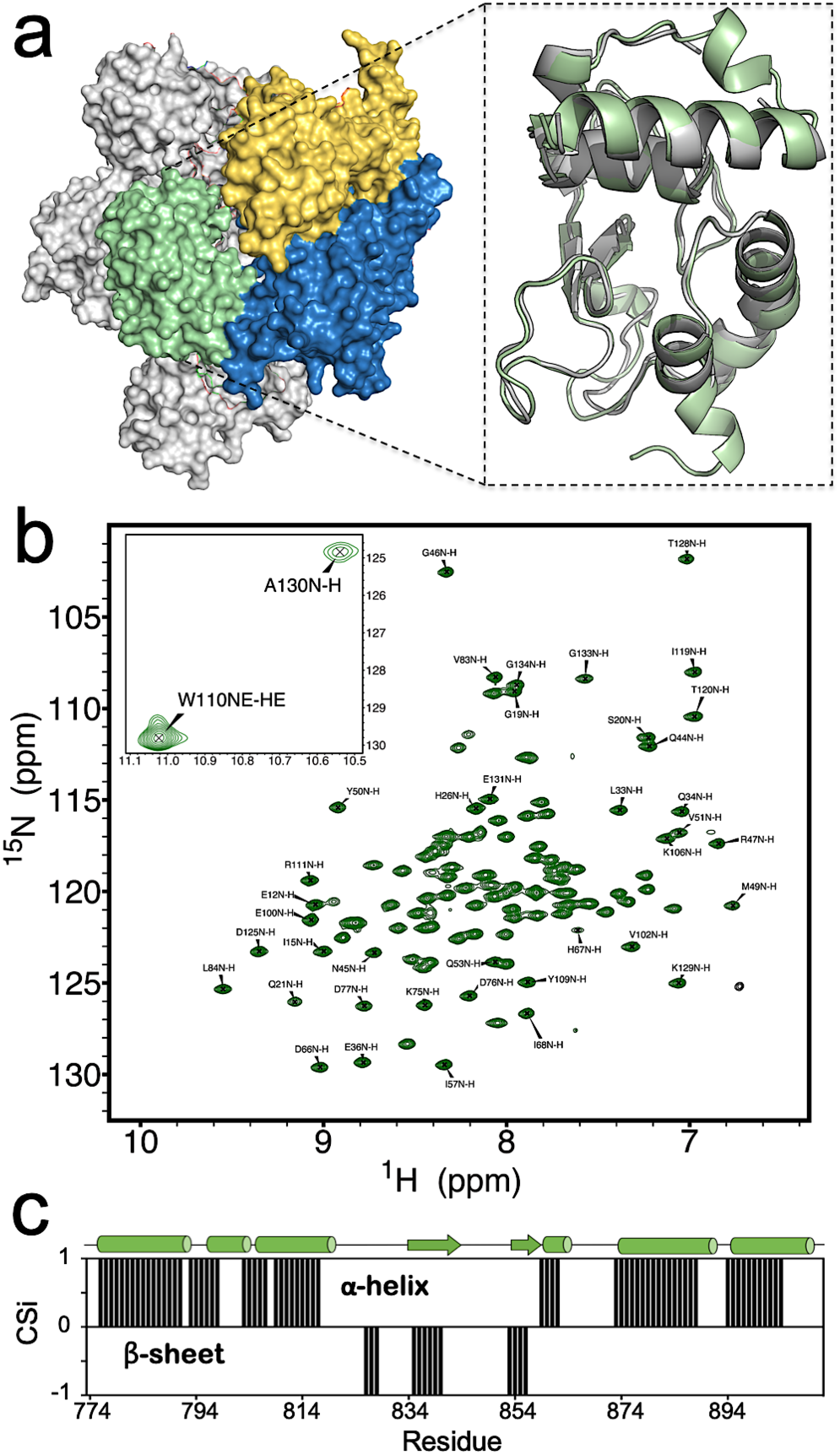
NMR Spectrum of HNH. **(a)** Architecture of the Cas9 protein (PDB code: 4UN3),^5^ highlighting its protein domains as follows: HNH (green), RuvC (blue), PAM interacting region (PI, gold) and recognition lobe (REC, gray). In the close-up view, a model of the HNH structure determined from NMR data (green) is overlaid with that of HNH from the full-length Cas9 (gray). (**b**) ^1^H-^15^N HSQC NMR spectrum of the HNH nuclease domain from S. pyogenes Cas9 (the inset reports two peaks out of range). (**c**) Consensus chemical shift index (CSi), indicating the predicted secondary structure for the HNH construct based on the NMR chemical shifts (black bars from 0 to 1 indicate α-helix, while bars from 0 to -1 indicate β-sheet, see the Methods section) compared to that of HNH from the full-length Cas9 (shown on top of the graph as sequence, with α-helical and β-sheet regions indicated as tubes and arrows).

Here, we probe the structural and dynamic determinants of allosteric signaling in the Cas9 HNH nuclease by means of solution NMR and all-atom MD simulations. A novel construct of the HNH nuclease domain from *S. pyogenes* Cas9 has been engineered that maintains the fold and properties of the wild-type (WT, i.e. full-length) Cas9 protein and allows the characterization of its multi-timescale conformational dynamics by solution NMR spectroscopy and MD simulations. To comprehensively access the slow timescale dynamics of the system at the atomic scale, we performed accelerated MD (aMD). aMD is an enhanced sampling methodology that applies a boost potential to the simulation, thereby accelerating transitions between low-energy states. The method has previously been shown to access slow dynamical motions in biomolecules, in excellent agreement with NMR experiments.^28-30^ However, the use of aMD for large biomolecular systems, such as CRISPR-Cas9, can suffer from high statistical noise, which hampers the characterization of the correct statistical ensemble.^31-32^ To overcome this limitation, a novel Gaussian accelerated MD (GaMD) method^33^ has been proposed, which uses harmonic functions to construct a boost potential that is adaptively added to the simulation (see the Methods section). GaMD has been applied to large biomolecular complexes, successfully describing the long timescale dynamics of CRISPR-Cas9 and G-protein coupled receptors. Hence, while overcoming the limitations of the early aMD methodology, GaMD still holds the method capability to describe slow dynamical motions, which are relevant for allostery,^19^ and to provide an atomic-level comparison with NMR experiments.^28-30^ As a result of our experimental and theoretical approach, we identify a dynamic pathway that connects HNH and RuvC through contiguous *ms* timescale motions, while also highlighting its propagation to the REC lobe to enable the information transfer for concerted cleavage of DNA. Structure-based prediction of the NMR chemical shifts further reveal the agreement between experiments and computations, indicating that the structural/dynamic features derived from GaMD simulations represent the experimental results well at the molecular level. Overall, the integrated approach employed in this study enabled access to the intrinsic conformational fluctuations of the Cas9 HNH nuclease, which are essential for allosteric signaling in CRISPR-Cas9. Our combined NMR and theoretical approach paves the way for the complete mapping of allosteric signaling and determination of its role in the enzymatic function and specificity of Cas9.

## Results

### Structural features of the HNH nuclease

To determine the structural features of the isolated HNH domain, we employed solution NMR and X-ray crystallography. First, the structure of the HNH domain (Figure 1A) was derived from the assigned ^1^H-^15^N HSQC NMR spectrum (Figure 1B) and backbone assignments.^34^ Backbone assignments were uploaded to the CS23D server in order to predict the HNH structure from composite NMR chemical shift indices.^35^ Figure 1A shows a model of the HNH structure determined from NMR data (close-up view, green) overlaid with that of HNH from the full-length Cas9 (gray). The predicted structure is remarkably similar to that of full-length Cas9 (PDB code: 4UN3)^5^ displaying Cα root-mean-squared-deviation RMSD = 0.688 Å. The NMR model also highlights small helical turns in regions of poor electron density in the full-length Cas9 structure, as well as an extension of the C-terminal α-helix. The secondary structure of this construct determined from Cα and Cβ chemical shift indices is in good agreement with that of the HNH domain from the full-length Cas9 (Figure 1C), indicating that the engineered protein is a good representation of this fold in solution. Circular dichroism (CD) spectroscopy is consistent with a predominantly α-helical protein (Figure S1), in agreement with the X-ray structure of the full-length Cas9.^5^ Interestingly, the thermal stability of this engineered construct is 12 °C higher than that of full-length Cas9, despite identical amino acid sequences, and is soluble to concentrations exceeding 1 mM.

The similarity of our construct to that of HNH from the fulllength Cas9 supports the reliability of the predicted structure. A further confirmation is provided by the X-ray crystal structure of the HNH construct solved at 1.3 Å resolution (Figure 2). This X-ray crystal structure aligns well to that of the full-length Cas9 (PDB code 4UN3)^36^ and the predicted NMR structure, with a Cα RMSD values of 0.549 Å and 0.479 Å, respectively, with the most significant difference due to a crystal contact in the experimental lattice pushing the N-terminal helix inward (Figure 2, inset top).

**Figure 2.**
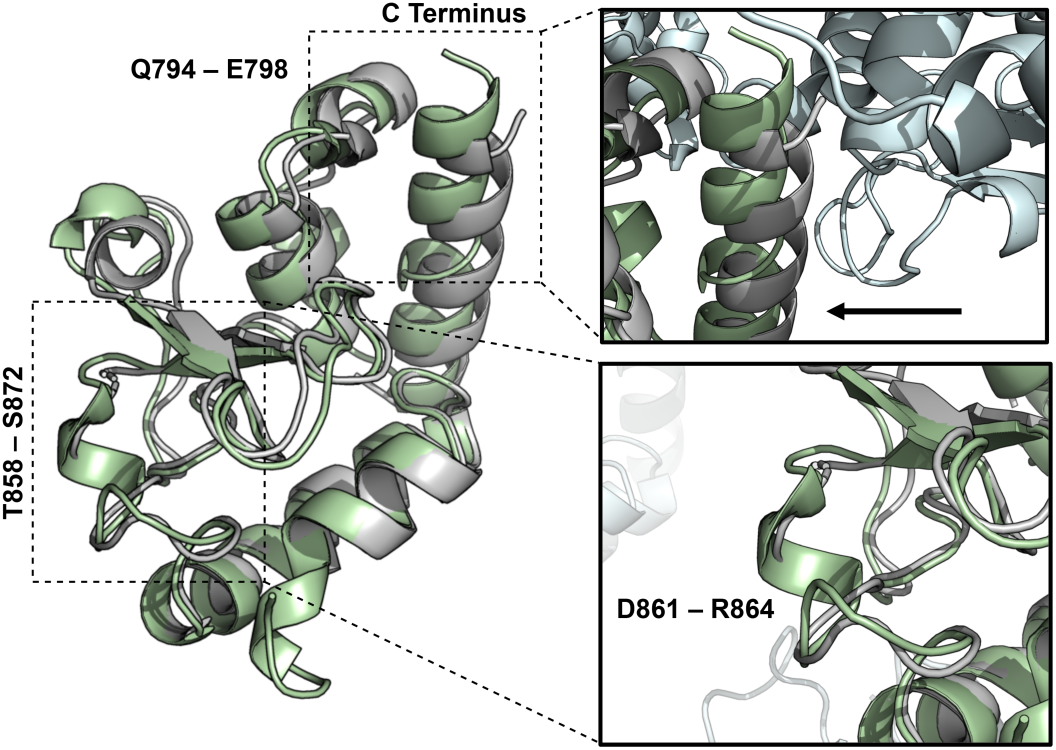
X-ray structure of HNH. The X-ray structure of the isolated HNH domain (PDB code: 6O56, green), solved at 1.30 Å resolution, is overlaid with the X-ray structure of the HNH domain from the full-length *S. pyogenes* Cas9 (PDB code: 4UN3, gray).^5^

The overall fold of HNH from full-length Cas9 is therefore well-maintained in the isolated domain. The residues L791–E802 and T858–S872 form two flexible loop regions, as suggested by NMR. An α-helix is introduced in residues Q794–E798 and an additional solvent exposed loop comprised of residues T858–S872 forms a small α-helix at D861–R864 (Figure 2, inset bottom), also observed in the structural model from the NMR chemical shifts. Lastly, a small extension of the C-terminal α-helix is also confirmed in this novel X-ray structure.

#### Experimental conformational dynamics of the HNH nuclease

In order to experimentally probe timescales relevant to allostery, we first analyzed the dynamics of HNH with the method of Bracken and coworkers.^37^ Here, potential sites of *ps*–*ns* and *μs*– *ms* flexibility have been identified through the analysis of the *R*_1_*R*_2_ product. Motions on each of these timescales contribute to allostery; *ps-ns* motions have been associated with allosteric activation as effector molecules induce favorable changes in configurational entropy,^38-39^ leading to a population shift from inactive-to-active states, while *μs*–*ms* processes are often commensurate with the rates of catalytic reactions.^40-41^ With respect to the individual longitudinal and transverse relaxation rates, the *R*_1_*R*_2_ product attenuates the contribution of motional anisotropy and more clearly illuminates sites of chemical exchange. The *R*_1_*R*_2_ values for each residue in HNH (Figure 3A and Table S1) highlight several locations of *ps*–*ns* and *μs*–*ms* flexibility. Twenty residues display *R*_1_*R*_2_ values above 1.5σ of the 10% trimmed mean, due to the significant influence of *R*_ex_ related to *μs*–*ms* motion. Measured *R*_ex_ parameters are consistent with this interpretation (Figures 3A, S3 and Table S1). A lower number of residues (i.e., 13) fall below 1.5σ of the mean, suggesting potential influence of *ps– ns* dynamics at these sites, with the mean *R*_1_*R*_2_ value corresponding to an average order parameter (*S*^2^) value of 0.85, where 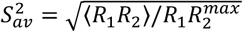. Steady-state ^1^H-[^15^N] NOE were also measured and the order parameter (*S*^2^) was determined for assigned residues in HNH with RELAX.^42^ Regions of *ps–ns* flexibility (*i.e.* high configurational entropy) are observed in residues 822– 843 and 890–904. Consistent with these data, in the X-ray structure of full-length Cas9 residues 822–843 are exposed toward the solvent, while residues 890–904 comprise flexible loop regions.^5^ In order to expand on this analysis, we quantified the conformational exchange parameters associated with millisecond dynamics of the HNH nuclease by Carr–Purcell–Meiboom–Gill (CPMG) relaxation dispersion experiments (Figure 3B and Table S2). Residues displaying slow timescale (*ms*) dynamics correspond to K782–E786, I788, K789, G792, Q794–E798, Y815-N818, R820, E827–D829, H840, V841, S851, D853, K855, E873 and L900 in the full-length Cas9. Many of these sites are also indicated by the *R*_1_*R*_2_ analysis. Rates of conformational exchange (k_ex_) at these sites range from 800 – 2900 s^-1^ with an average k_ex_ = 1761 ± 414. Interestingly, k_ex_ values determined from CPMG experiments show an approximate bimodal distribution, with a large number of residues showing 1000 ≤ k_ex_ ≤ 2000 s^-1^ and fewer residues with 2000 ≤ k_ex_ ≤ 3000 s^-1^ (Figure S4). Regions with larger k_ex_ values are primarily confined to the periphery and N-terminus of HNH. Residues with smaller k_ex_ values comprise the majority of the HNH-REC interface and central dynamic core.

**Figure 3.**
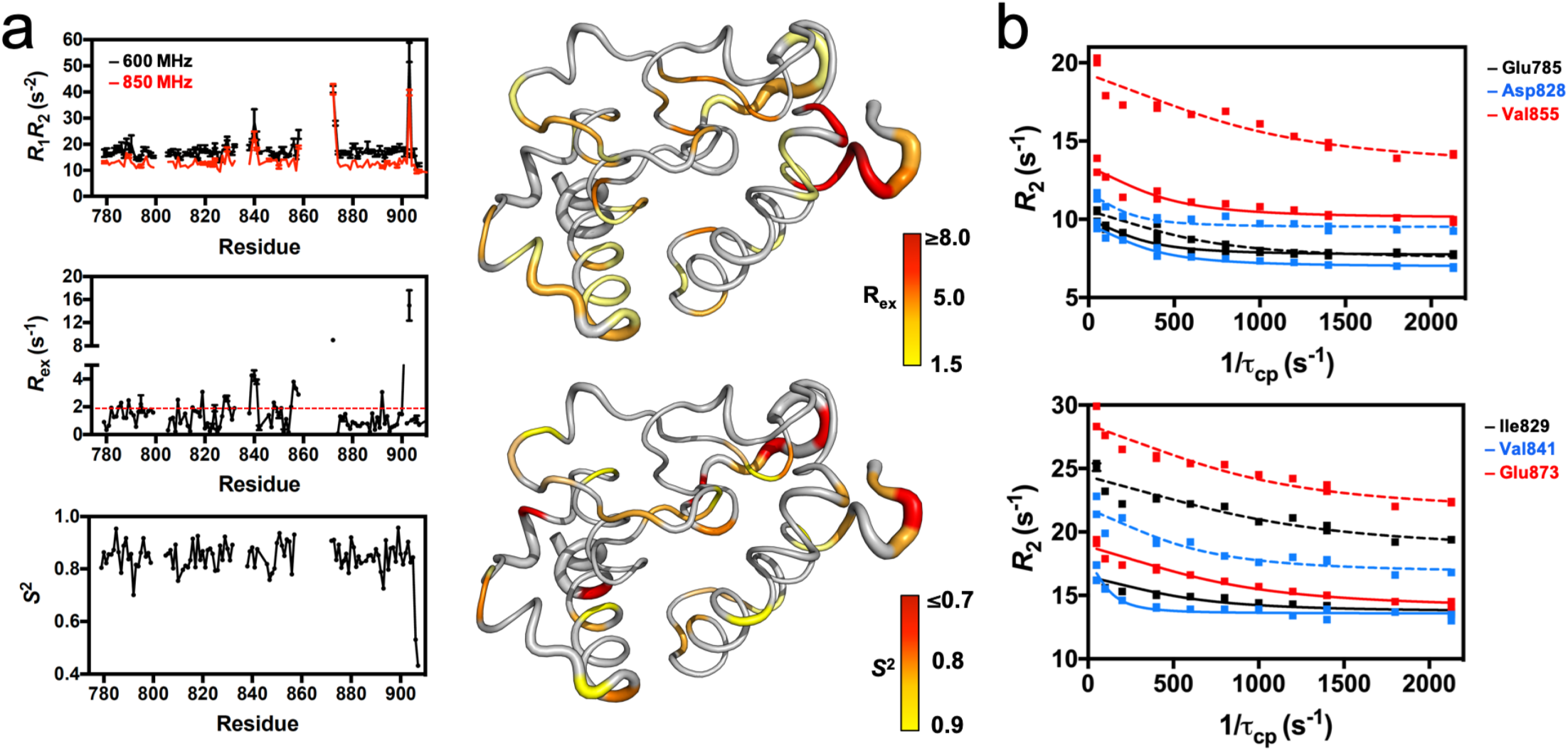
HNH Dynamics Measured by NMR. (**a**) Plots of *R*_1_*R*_2_, *R*_ex_, and the order parameter (*S*^2^, determined from model-free analysis of *T*_1_, *T*_2_, ^1^H-[^15^N]-NOE measurements) for the HNH nuclease. The *R*_1_*R*_2_ parameters were measured at 600 (black) and 850 (red) MHz. The red dashed line on the *R*_ex_ plot denotes the 1.5σ from the 10% trimmed mean of the data. The *R*_ex_ (right, top) and *S*^2^ (right, bottom) values are mapped onto the HNH structure and colored according to the adjacent legends. (**b**) Selected CPMG relaxation dispersion curves collected at 600 (solid lines) and 850 (dashed lines) MHz.

### Allosteric signaling pathway

The *ms* dynamics identified via CPMG relaxation dispersion experiments are of particular interest for the identification of the Cas9 signaling pathways, due to the established importance of slow dynamical motions in allosteric regulation.^19^ Residues displaying slow timescale (*ms*) dynamics (Table S2) form clusters in three regions of HNH (Figure 4, red spheres), two of which are the interface with the region REC2 of the recognition lobe and with RuvC (i.e., the HNH–REC2 and HNH–RuvC interfaces), while the third region is located in the core of HNH. This well-defined subset of flexible residues within HNH therefore bridges the RuvC and REC2 interfaces, forming a contiguous dynamic pathway within the isolated HNH domain. This pathway of flexible residues connecting HNH–RuvC and HNH–REC2 agrees well with the available experimental evidence that indicate the existence of allosteric communication within CRISPR-Cas9. A tight dynamic inter-connection between HNH and RuvC has been originally reported by Sternberg and colleagues^9^ and supported by MD simulations studies.^17^ Moreover, the REC2 region has been recently suggested to be involved in the activation of HNH through an allosteric regulation that also implicates the REC3 region.^36^ The authors have shown that, upon binding a complementary RNA:DNA structure prone to undergo DNA cleavage, the REC3 region modulates the motions of the neighboring REC2, which in turn contacts HNH and sterically regulates its access to the scissile phosphate. MD simulations of the fully activated CRISPR-Cas9 complex revealed that highly coupled motions between REC2, REC3 and HNH are critical for the activation of the catalytic domain toward cleavage, supporting the existence of an allosteric signal.^43^ A recent experimental study has further suggested that REC2 is critical in regulating the rearrangements of the DNA for double strand cleavage via the HNH and RuvC nuclease domains.^44^ Taken together, these findings strongly support the outcomes of the NMR experiments reported here, suggesting that the dynamic pathway spanning the isolated HNH domain is responsible for the information transfer between RuvC and REC2.

**Figure 4.**
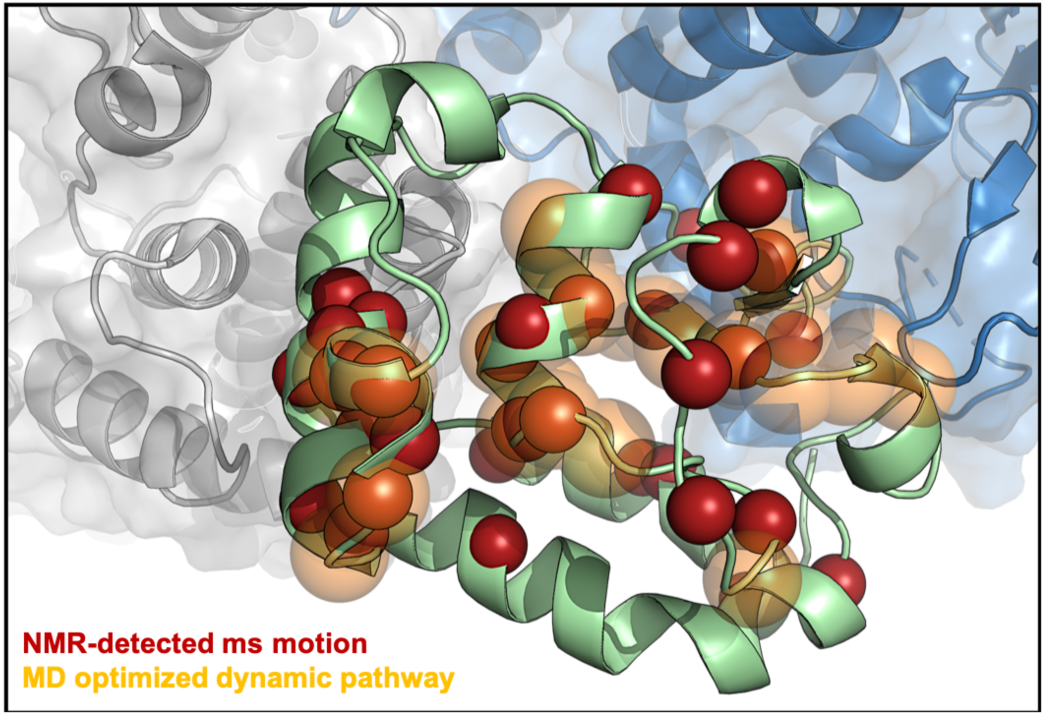
Allosteric signaling across HNH. Flexible residues in the HNH construct (green) measured by CPMG relaxation dispersion NMR (red spheres) and the theoretical allosteric pathway (orange, transparent) optimizing the overall correlation between HNH residues at the RuvC (blue) and REC2 (gray) interfaces. A significant number of the sites identified by NMR are within the top 10 optimal pathways calculated from MD trajectories, suggesting that the experimentally measured dynamic pathway spanning the HNH domain correlates well with the computationally derived pathway for optimal information transfer.

To gain insight into the allosteric signaling pathway within the full-length Cas9,^5^ the protein was subjected to extensive computational analyses that are suitable for the detection of allosteric analyses that are suitable for the detection of allosteric effects.^41, 45-47^We combined correlation analyses and network models derived from graph theory to determine the most relevant pathways across HNH communicating RuvC with REC2. The computed pathways are composed by residue–to–residue steps that optimize the overall correlation (i.e., the momentum transport) between amino acids 789/794 and 841/ 858 (belonging to HNH but adjacent to RuvC and REC2, respectively). This yields an estimation of the principal channels of communication between RuvC and REC2. Interestingly, the pathway that maximizes the dynamic transmission between RuvC and REC2 through HNH (Figure 5, orange spheres) agrees remarkably well with the pathway experimentally identified in the HNH construct via CPMG relaxation dispersion (Figure 5, red spheres). Residues belonging to the computational pathway are G792*, Q794*, K797*, E798*, Y812, L813, Y814, L816*, Q817*, N818*, G819, R820*, D825, I830, V838*, D839, H840*, I841*, V842, P843†, Q844†, N854, K855*, V856, L857, T858, R859†, S860†, D861†,K862†; where the asterisk indicates that they also show slow dynamics in the HNH construct (as experimentally identified via CPMG relaxation dispersion and *R*_1_*R*_2_ (+1.5σ), Tables S1-S2) and the dagger indicates residues unassigned by NMR. Importantly, while our experimental approach is restricted to the slow dynamic residues, the computation of the optimal pathways considers motions in every possible time-scale within the simulated window. Therefore, differences between the two approaches may be originated in the fast-dynamic component present in the simulated pathways. On the other hand, several differences between the relaxation dispersion network and computationally derived network are due to missing NMR assignments for these residues (daggers). These missing assignments are, in some cases, due to exchange broadening of the NMR signal, indicative of a highly flexible site. Also of note, are the many sites of NMR-detected *ms* motions directly adjacent to the MD optimized pathway. Examples from the MD (NMR) pathways are 1) Y812 – Y814 (Y815); 2) D825, I830 (E827, L828, D829); 3) D839 (V838, H840); and 4) N854, V856, L857, T858 (D853, K855). This consensus between the dynamic pathways experimentally observed in the HNH construct by NMR and in full-length Cas9 by MD indicates that the REC2–HNH–RuvC communication channel is conserved in the full-length enzyme.

**Figure 5.**
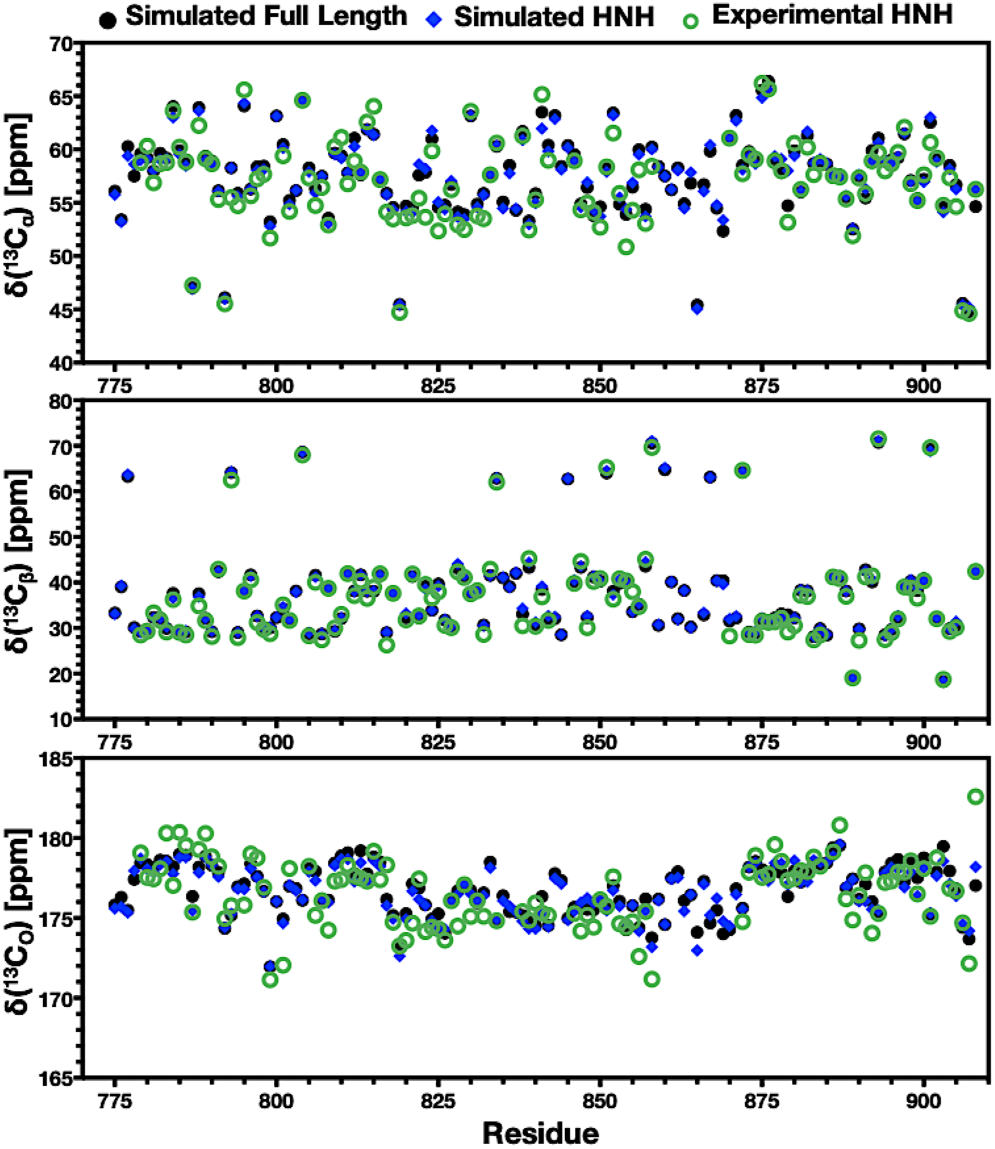
Experimental vs. simulated carbon chemical shifts. Experimental and simulated NMR chemical shifts of ^13^C_α_ (top), ^13^C_β_ (center), and ^13^C_O_ (bottom) plotted for each residue in the HNH domain. Data in each plot compare the experimental chemical shifts (green circles) with those calculated from simulations of the isolated HNH domain (blue diamonds) and of the full-length Cas9 complex (black circles). All simulated spectra were computed as described in Methods utilizing GaMD trajectories.

### Conformational dynamics of HNH in full-length Cas9

In order to compare the conformational dynamics of this novel HNH construct with those of the full-length Cas9, and to further interpret the outcomes of solution NMR experiments, we performed all-atom MD simulations on the structure of the HNH domain predicted by NMR and the X-ray structure of the fulllength Cas9.^5^ To access the slower timescale dynamics of the systems, we performed accelerated MD simulations, using a Gaussian accelerated MD (GaMD) method,^33^ which has been shown to describe the *µs* and *ms* dynamics of CRISPR-Cas9 quite well.^14, 48-49^ Indeed, while classical MD can detect fast stochastic motions responsible for spin relaxation, more sophisticated methods that enhance the sampling of the configurational space are required to access the slower motion quantified by solution NMR. Accelerated MD is a biased-potential method,^50^ which adds a boost potential to the potential energy surface (PES), effectively decreasing the energy barriers separating low-energy states, thus accelerating the occurrence of slower dynamic events. As shown by several independent reports, the method accurately reproduces the slow dynamics captured by solution NMR in biomolecular systems,^28-30^ therefore providing comparison with the experimental results reported here.

The simulated trajectories have been analyzed to compare the conformational dynamics of HNH in its isolated form and in fulllength Cas9. By performing Principal Component Analysis (PCA), the dynamics of HNH along the first principal mode of motion – usually referred as *“essential dynamics”*^51^ – reveals remarkable similarities in the full-length Cas9 and in the isolated form (Figures S5-S6). Interestingly, the residues of HNH that experimentally display *ms* dynamics (i.e., as captured from the CPMG relaxation dispersion and *R*_1_*R*_2_ (+1.5σ) measurements) are characterized by short amplitude motions in both the isolated form of HNH and when embedded in the full-length Cas9. Analysis of the root mean square fluctuations (RMSF) of individual Cα atoms further shows that the residues with slow timescale motions (experimentally identified via NMR) display small fluctuations in the simulations of both the isolated HNH and in the full-length Cas9 (Figure S7). This indicates that short amplitude motions and small fluctuations are conserved in the regions that form a continuous *ms* dynamic pathway connecting REC2–HNH–RuvC (Figure 4). In this respect, it is important to note that short amplitude motions, as well as small fluctuations, do not directly correspond to slow time scale dynamical motions. However, the consensus observed in both the HNH construct and within the full-length Cas9 indicates similar intrinsic dynamics along the pathway connecting REC2–HNH–RuvC, which has been experimentally derived via NMR (Figure 4). Inspection of the conformational ensemble accessed during the simulations reveals that the isolated HNH domain resembles the ensemble of the full-length system overall, with a remarkable similarity in terms of short amplitude motions and low fluctuations for the residues within the REC2–HNH–RuvC pathway (Figure S8). Overall, the analysis of the conformational dynamics shows that the HNH construct maintains the fold observed in full-length Cas9, supporting the connection between conformational dynamics captured via solution NMR and those of HNH inside full-length Cas9.

### Simulated ensemble and NMR experiments

To gain insight into how well the structural and dynamical features captured by GaMD simulations represent the NMR experiments at the molecular level, the simulated trajectories were used to compute NMR chemical shifts with the SHIFTX2 code.^52^ We detected excellent sequence-specific agreement between predicted and experimental C_α_, C_β_, and C_O_ chemical shifts for the isolated HNH domain (Figure 5). The experimental distributions of backbone amide groups also display remarkable agreement between experimental and simulated values (Figure S9) This is a strong indication that the GaMD ensemble properly represents the NMR experiments at a molecular level. Another important aspect of these simulations is the similarity of HNH in full-length Cas9 and its isolated form. Figure 5 shows that these forms of HNH display very similar C_α_, C_β_, and C_O_ chemical shifts as well, indicating that HNH presents similar spectral trends when it is isolated or in full-length Cas9. This is therefore a further indication that the structural dynamics of the HNH construct predicted by NMR are comparable to those of HNH in full-length Cas9, supporting the comparison performed here. Importantly, the observed agreement between the computed and experimental spectra is also observed in the simulation replicas (Figure S9).

## Discussion

The power of the CRISPR-Cas9 system is its ability to perform targeted genome editing *in vivo* with high efficiency and increasingly improved specificity.^36, 53-55^ In this system, an intriguing allosteric communication has been suggested to propagate the DNA binding signal across the HNH and RuvC nuclease domains to facilitate their concerted cleavage of the two DNA strands.^9, 17^ In this process, the intrinsic flexibility of the catalytic HNH domain would regulate the information transfer, exerting conformational control. Here, solution NMR experiments are used to capture the intrinsic motions responsible for the allosteric signaling across the HNH domain. We use a novel engineered construct of HNH that maintains the conformation and flexibility of full-length Cas9 to reveal the existence of a *ms* timescale dynamic pathway. This network spans the HNH domain from the region interfacing the RuvC domain and propagates up to the REC lobe at the level of the REC2 region (Figure 4). In-depth analysis of the allosteric signaling within the full-length Cas9 has been performed, by employing theoretical approaches that are suited for the detection of allosteric effects.^41, 45-47^ Indeed, the dynamic pathway experimentally observed in the HNH construct is conserved in the fulllength Cas9 (Figure 4), confirming the existence of a communication channel between REC2–HNH–RuvC. This continuous pathway confirms the direct communication between the two catalytic domains, originally identified by the experimental work of Sternberg^9^ and supported by MD simulations,^17^ and also discloses their connection to the REC2 region. In this respect, single molecule FRET experiments have indicated that REC2 is critical for the activation of HNH through an allosteric mechanism that also involves the REC3 region.^36^ Accordingly, in the fully activated complex, the REC3 region would modulate the motions of the neighboring REC2, which in turn contacts HNH and regulates its access to the scissile phosphate. By doing so, the REC region would act as a *“sensor”* for the formation of a RNA:DNA structure prone to DNA cleavage, transferring the DNA binding information to the catalytic HNH domain in an allosteric manner. A tight dynamical interplay between REC2–REC3 and HNH has also been detected via MD simulations of the fully activated CRISPR-Cas9 complex, revealing that highly coupled motions of these regions are at the basis of the activation of HNH for DNA cleavage.^43^ A recent important contribution further suggested that REC2 regulates the rearrangements of the DNA to attain double strand cleavages via the HNH and RuvC nucleases.^44^ Altogether, these experimental outcomes strongly support the finding of a continuous dynamic pathway spanning HNH from RuvC to REC2, and suggest its functional role for the allosteric transmission. To further investigate the motions associated with allosteric signaling, the conformational dynamics of the HNH domain was investigated by means of accelerated MD simulations, which can probe long-timescale *µs* and *ms* motions in remarkable agreement with NMR experiments.^28-30^ Analysis of these conformational dynamics indicates that the HNH construct maintains the overall fold observed in full-length Cas9, and indicates conserved short amplitude motions and low fluctuations in the regions that form a continuous *ms* dynamic pathway connecting REC2–HNH–RuvC. Taken together, these computational outcomes suggest that the intrinsic conformational dynamics experimentally identified in the HNH construct reasonably resemble the dynamics of HNH in the full complex, supporting the connection between the two systems. Finally, mixed machine learning and structure-based prediction of the NMR chemical shifts from the simulated trajectories have also revealed the agreement between experiments and computations, indicating that the structural/dynamic features derived via GaMD simulations represent the experimental results at the atomic and molecular level.

Overall, by combining solution NMR experiments and MD simulations, we identified the dynamic pathway for information transfer across the catalytic HNH domain of the CRISPR-Cas9 system. This pathway, which spans HNH from the RuvC nuclease interface up to the REC2 region in the full-length Cas9, is suggested to be critical for allosteric transmission, propagating the DNA binding signal across the recognition lobe and the nuclease domains (HNH and RuvC) for concerted cleavage of the two DNA strands. This study also represents the first step toward a complete mapping of the allosteric pathway in Cas9 through solution NMR experiments. In this respect, despite modern experimental practices such as perdeuteration,^56^ transverse relaxation-optimized spectroscopy (TROSY),^57^ sparse isotopic labeling,^58^ and ^15^N-detection,^59^ the complete characterization of the slow dynamical motions responsible for the allosteric signaling has remained challenging, due to the size of the polypeptide chain of the Cas9 protein (∼ 160 kDa). Future investigations – reliant upon ongoing experiments and computations in our research groups – will include the investigation of the information transfer between HNH and RuvC and the allosteric role of their flexible interconnecting loops.^9, 17^ Further, our joint NMR/MD investigations are being employed to understand the role of the recognition region within the allosteric activation. This is of key importance, since mutations within the REC lobe – at distal sites with respect to HNH – can control the activation of HNH and the specificity of the enzyme toward on-target DNA sequences.^36, 53-55^ As such, by providing fundamental understanding of the intrinsic allosteric signaling within the catalytic HNH domain, the present study poses the basis for the complete mapping of the allosteric pathway in Cas9 and its role in the on-target specificity, helping engineering efforts aimed at improving the genome editing capability of the Cas9 enzyme.

## Materials and Methods

### Protein Expression and Purification

The HNH domain of *S. pyogenes* Cas9 (residues 775-908) was engineered into a pET15b vector with an N-terminal His_6_-tag and expressed in Rosetta(DE3) cells in M9 minimal medium containing MEM vitamins, MgSO_4_ and CaCl_2_. Cells were induced with 0.5 mM IPTG after reaching an OD_600_ of 0.8 – 1.0 and grown for 16 – 18 hours at 22 ºC post induction. The cells were harvested by centrifugation, resuspended in a buffer containing 20 mM HEPES, 500 mM KCl, and 5 mM imidazole at pH 8.0, lysed by ultrasonication and purified on a Ni-NTA column. NMR samples were dialyzed into a buffer containing 20 mM HEPES, 80 mM KCl, 1 mM DTT and 7.5% (v/v) D_2_O at pH 7.4.

### X-ray Crystallography

Following TEV cleavage of the His_6_-tag, HNH was subsequently purified by HiPrep 16/60 Sephacryl 100 S-100 HR gel filtration chromatography. Crystals were obtained with sitting drop vapor diffusion at room temperature with 48 mg/mL HNH 1:1 with the Molecular Dimensions Morpheus I Screen condition E4 (0.1 M mixture of [imidazole and MES] pH 6.5, 25% (v/v) mixture of [2-methyl-2,4-pentanediol, PEG1000, and PEG3350], and 0.3 M mixture of [diethylene glycol, triethylene glycol, tetraethylene glycol, and pentaethlyene glycol]). Diffraction data were collected on a Rigaku MicroMax-003i sealed tube X-ray generator with a Saturn 944 HG CCD detector and processed and scaled using XDS^60^ and Aimless in the CCP4 program suite.^61^ The HNH domain from full-length *S. pyogenes* Cas9 was used for molecular replacement (PDB: 4UN3)^5^ with Phaser in the PHENIX software package.^62^ Iterative rounds of manual building in Coot^63^ and refinement in PHENIX yielded the final HNH domain structure.

### NMR Spectroscopy

NMR spin relaxation experiments were carried out at 600 and 850 MHz on Bruker Avance NEO and Avance III HD spectrometers, respectively. All NMR spectra were processed with NMRPipe ^64^ and analyzed in SPARKY.^65^ Backbone chemical shift data was uploaded to the CS23D server for secondary structure calculations. Carr-Purcell-Meiboom-Gill (CPMG) NMR experiments were adapted from the report of Palmer and coworkers,^66^ and performed at 25 ºC with a constant relaxation period of 40 ms, a 2.0 second recycle delay, and τ_cp_ points of 0.555, 0.625, 0.714, 0.833, 1.0, 1.25, 1.5, 1.667, 2.5, 5, 10, and 20 ms. Relaxation dispersion profiles were generated by plotting *R*_2_ vs. 1/τ_cp_ and exchange parameters were obtained from fits of these data carried out with in-house scripts and in RELAX under the R2eff, NoRex, Tollinger (TSMFK01), and Carver-Richards (CR72 and CR72-Full) models.^42, 67^ Two-field relaxation dispersion data were fit simultaneously and uncertainty values were obtained from replicate spectra (see the Supporting Information, SI). Longitudinal and transverse relaxation rates were measured with relaxation times of 0(x2), 40, 80, 120, 160(x2), 200, 240, 280(x2), 320, 360, and 400 ms for <u>T_1_</u> and 4.18, 8.36(x2), 12.54, 16.72, 20.9(x2), 25.08(x2), 29.26, 33.44, 37.62, and 41.8 ms for <u>T_2_</u>. Peak intensities were quantified in Sparky and the resulting decay profiles were analyzed in Mathematica with errors determined from the fitted parameters. Steady-state ^1^H-[^15^N] NOE were measured with a 6 second relaxation delay followed by a 3 second saturation (delay) for the saturated (unsaturated) experiments. All relaxation experiments were carried out in a temperature-compensated interleaved manner. Model-free analysis using the Lipari-Szabo formalism was carried out on dual-field NMR data in RELAX with fully automated protocols.^42^

### Computational Structural Models

Two model systems were built for MD simulations, the first of which was based on the X-ray structure of the full-length wild-type Cas9 protein in complex with RNA and DNA, solved at 2.58 Å resolution (PDB code: 4UN3).^5^ The second model system was based on the NMR structure of the HNH domain obtained in this work. The RMSD between the HNH domain in the X-ray structure of the WT Cas9 complex^5^ and the HNH domain structure determined here is 0.688 Å. Both model systems were embedded in explicit water, adding Na+ counter-ions to neutralize the total charge, reaching a total of ∼220,000 atoms and a box size of ∼145 × 110 × 147 Å^3^ for the CRISPR-Cas9 complex and ∼25,000 atoms and a box size of ∼72 × 62 × 60 Å^3^ for the HNH domain.

### Molecular Dynamics (MD) Simulations

The above-mentioned model systems were equilibrated through conventional MD. We employed the Amber ff12SB force field, which includes the ff99bsc0 corrections for DNA^68^ and the ff99bsc0+*χOL3* corrections for RNA.^69-70^ Hydrogen atoms were added assuming standard bond lengths and constrained to their equilibrium position with the SHAKE algorithm. Temperature control (300 K) was performed via Langevin dynamics,^71^ with a collision frequency γ = 1. Pressure control was accomplished by coupling the system to a Berendsen barostat,^72^ at a reference pressure of 1 atm and a relaxation time of 2 ps. All simulations have been carried out through a well-established protocol described in the SI. MD simulations were carried out in the NVT ensemble, collecting ∼100 ns for each system (for a total of ∼400 ns of production runs). These well-equilibrated systems have been used as the starting point for Gaussian accelerated MD (GaMD, details below). All simulations were performed with the GPU version of AMBER 16.^73^

### Gaussian Accelerated MD Simulations (GaMD)

Accelerated MD (aMD) is an enhanced sampling method that adds a boost potential to the Potential Energy Surface (PES), effectively decreasing the energy barriers and accelerating transitions between low-energy states.^50^ The method extends the capability of MD simulations over long timescales, capturing slow *µs* and *ms* motions in excellent comparability with solution NMR experiments.^28-30^ Here, we applied a novel and robust aMD method, namely a Gaussian aMD (GaMD),^33^ which uses harmonic functions to construct a boost potential that is adaptively added to the PES, enabling unconstrained enhanced sampling and simultaneous reweighting of the canonical ensemble.

Considering a system with *N* atoms at positions 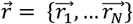, when the system potential 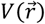 is lower than a threshold energy *E*, the energy surface is modified by a boost potential as:

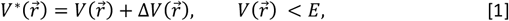

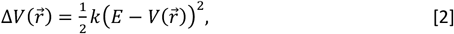

where *k* is the harmonic force constant. The two adjustable parameters *E* and *k* are automatically determined by applying the following three criteria. First, for any two arbitrary potential values 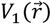 and 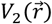 found on the original energy surface, if 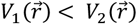, Δ*V* should be a monotonic function that does not change the relative order of the biased potential values, *i.e*. 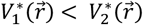. Second, if 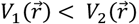, the potential difference observed on the smoothed energy surface should be smaller than that of the original, *i.e.* 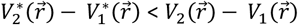. By combining the first two criteria with Eqn [1] and [2]:

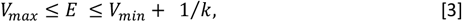

where *V*_*min*_ and *V*_*max*_ are the system minimum and maximum potential energies. To ensure that Eqn. [4] is valid, *k* must satisfy *k* ≤ 1/*V*_*max*_ − *V*_*min*_. By defining *k* ≡ *k*_0_ 1/*V*_*max*_ − *V*_*min*_, then 0 < *k* ≤ 1. Lastly, the standard deviation of Δ*V* must be narrow enough to ensure accurate reweighting using cumulant expansion to the second order: *σ*_Δ*V*_ = *k*(*E* − *V*_*avg*_)*σ*_*V*_ ≤ *σ*_0_, where *V*_*avg*_ and *σ*_*V*_ are the average and standard deviation of the system potential energies, *σ*_Δ*V*_ is the standard deviation of Δ*V* and *σ*_0_ as a user-specified upper limit (e.g., 10 *k*_*B*_T) for accurate reweighting. When *E* is set to the lower bound, *E* = *V*_*min*_, according to Eqn. [4], *k*_0_ can be calculated as:

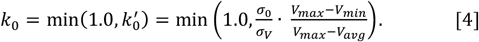

Alternatively, when the threshold energy *E* is set to its upper bound *E* = *V*_*min*_ + 1/*k, k*_0_ is:

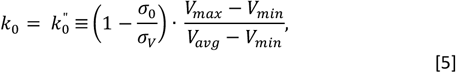

if 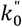 is calculated between *0* and *1*. Otherwise, *k*_0_ is calculated using Eqn. [4], instead of being set to 1 directly as described in the original paper.^33^ GaMD yields a canonical average of an ensemble by reweighting each point in the configuration space on the modified potential by the strength of the Boltzmann factor of the bias energy, *exp* [*β*Δ*V*(*r*_*t*(*i*)_)] at that particular point.

Based on extensive tests on the CRISPR-Cas9 system,^14, 48-49^ the system threshold energy is *E* = *V*_*max*_ for all GaMD simulations. The boost potential was applied in a *dual-boost* scheme, in which two acceleration potentials are applied simultaneously to the system: *(i)* the torsional terms only and *(ii)* across the entire potential. A timestep of 2 fs was used. The maximum, minimum, average, and standard deviation values of the system potential (*V*_*max*_, *V*_*min*_, *V*_*avg*_ and *σ*_*V*_) were obtained from an initial ∼12 ns NPT simulation with no boost potential. GaMD simulations were applied to the CRISPR-Cas9 complex and our HNH domain construct. Each GaMD simulation proceeded with a ∼50 ns run, in which the boost potential was updated every 1.6 ns, thus reaching equilibrium. Finally, ∼400 ns of GaMD simulations were carried in the NVT *ensemble* out for each system in two replicas, for a total of ∼1.6 μs of GaMD. This simulation length (i.e., ∼400 ns per replica) has shown to exhaustively explore the conformational space of the CRISPR-Cas9 system.^14, 48^

### Determination of the Allosteric Pathways across the HNH domain

The allosteric pathway for information transfer has been investigated by employing correlation analysis and graph theory.^41, 45-47^ First, the generalized correlations (GC_*ij*_), which capture non-collinear correlations between pairs of residues *i* and *j*, are computed (details are in the SI).^74^ In a second phase, the GC_*ij*_ are used as a metric to build a dynamical network model of the protein.^47^ In this model, the protein amino acids residues constitute the nodes of the dynamical network graph, connected by edges (residue pair connection). Edge lengths, i.e., the inter-node distances in the graph, are defined using the *GC*_*ij*_ coefficients according to:

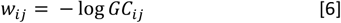

In the present work, two nodes have been considered connected if any heavy atom of the two residues is within 5 A° of each other (i.e., *distance cutoff*) for at least the 70 % of the simulation time (i.e., *frame cutoff*). This leads to the definition of a set of elements *w*_*ij*_ of the graph. In the third phase of the protocol, the optimal pathways for the information transfer between two nodes (i.e., two amino acids) are defined using the Dijkstra algorithm, which finds the roads, composed by inter-node connections, that minimize the total distance (and therefore maximize the correlation) between amino acids. In the present study, this protocol was applied on the trajectories of the full-length Cas9 simulated for ∼400 ns of GaMD simulations and averaged over two replicas. The Dijkstra algorithm was applied between the amino acids 789 and 841, which belong to HNH and are located at the interface with RuvC and REC2, respectively. As a result, the routes that maximize the correlation between amino acids 789 and 841 are identified, providing residue–to–residue pathways that optimize the correlations (i.e., the momentum transport). With the optimal motion transmission pathway the following 50 *sub-optimal* information channels where computed and accumulated and plotted on the 3D structure (Figure 4), to account for the contribution of the most likely sub-optimal pathways.

## Supporting Information

**Experimental procedures, circular dichroism spectra, NMR relaxation parameters, principal component analysis, MD simulations, and simulated NMR chemical shift distributions are available free-of-charge at pubs.acs.org.**

## Notes

<u>The authors declare no competing financial interest.</u>

## Acknowledgments

We thank Prof. Martin Jinek for the WT Cas9 plasmid and Prof. Patrick Loria for helpful discussion of spin relaxation. We thank Prof. Luca Mollica for useful discussions. This work was supported by start-up funds from Brown University and funds from the COBRE Center for Computational Biology of Human Disease (P20GM109035) to GPL, and by start-up funds from the University of California, Riverside to GP. Computer time for MD has been awarded by XSEDE via grant TG-MCB160059 (to GP).

